# A LAMP-based microfluidic chip for rapid detection of pathogen in Cryptococcal meningitis

**DOI:** 10.1101/2020.11.16.386045

**Authors:** Yueru Tian, Tong Zhang, Jian Guo, Huijun Lu, Yuhan Yao, Xi Chen, Xinlian Zhang, Guodong Sui, Ming Guan

## Abstract

Cryptococcal meningitis (CM) is a global threat with significant attributable morbidity and mortality. Information on integrated detection for CM diagnosis is still limited. This is mainly due to the presence of a large polysaccharide capsule and the tough cell wall of *Cryptococcus*, which makes it difficult to extract nucleic acids on the chip. In this study, we developed a LAMP-based microfluidic chip for rapid detection of pathogen in CM. We adopted 4 duplicate filtration membrane structures to improve target capture and simplify the enrichment process, and combined lyticase digestion and thermal alkaline lysisto optimize the nucleic acid extraction of *Cryptococcus* on the chip, and selected a portable UVA flashlight to shine the LAMP products to obtain the visual detection results which could be observed by the naked eye. This microfluidic chip, integrating sample *Cryptococcus* enrichment, nucleic acid extraction and LAMP detection unit, streamlined the operation process and reduced the exposure risk of directly handling cryptococcal samples. It did not require any additional instruments and demonstrated a rapid, reliable, as well as high-efficiency approach. It truly realized the “sample-to-answer” application and could be easily used for clinical cryptococcal prediagnosis.

## Introduction

*Cryptococcus* is an opportunistic pathogenic fungus that resides in diverse ecological niches, especially abundant in eucalyptus, avian excreta and amoeba. Among the 70 species identified, *C. neoformans* and *C. gatti* are the major causative agents of human cryptococcosis[1]. Found widely in soil, decaying wood, and bird droppings, *Cryptococcus* spores floating in the environment are inhaled by humans, causing pneumonia in immunocompromised patients, but in immunocompetent hosts the fungal cells are either cleared by the immune system or establish an asymptomatic latent infection. Once the infected subject becomes immunosuppressed, the latent infection can disseminate to other tissues, most notably the central nervous system (CNS), causing life-threatening subacute meningitis[1].

Cryptococcal meningitis (CM) is the most common cause of HIV-related meningitis, causing an estimated 223 100 (95% CI 150 600-282 400) incident cases and 181 100 (95% CI 119 400-234 300) deaths per year[2]. Globally, CM was responsible for 15% of HIV-related deaths[2]. In recent years, an increasing number of CM cases have been reported in non-HIV infected individuals, mainly including patients with natural and iatrogenic immunosuppression, such as patients with organ or stem-cell transplantation, malignant tumors, autoimmune diseases, glucocorticoid or immunosuppressive therapies (including chemo-, radio-, or immunotherapy for cancer), or even “immunocompetent individuals” carrying underlying immune deficiencies, chronic liver, kidney or lung disease and diabetes[3]. In-hospital mortality of non-HIV-related CM has now reached approximately 25% [4,5]. However, there are still about 65–70% of non-HIV CM patients without any predisposing factors, particularly in China[6-8], Korea[9], Japan[10] and India[11]. Moreover, *C. gattii* has been considered as the culprit causing CM in immunocompetent hosts, highlighted in Australia, Canada, and the U.S. Pacific Northwest[12]. Worldwide, the mortality rate of patients with *C. gattii* infections ranges from 13% to 33%[12].

Currently, the diagnosis of CM still poses some difficulties. One is attributed to atypical symptoms (fever, headache, nausea, vomiting), which are easily confused with upper respiratory tract infection, tuberculous meningitis, viral meningitis or neurological diseases. Another is that conventional CM diagnostic techniques have some limitations. Both of these challenges lead to delayed diagnosis and higher morbidity and mortality. To date, diagnosis of CM has mainly relied on cerebrospinal fluid (CSF) India ink microscopic staining, culture, cryptococcal anhtigen (CrAg) testing or histopathological examination. Although india ink microscopic staining has been a quick method, unfortunately sensitivity is low (86% in expert hands)[13]. The cultivation technique is time-consuming and usually takes 3-7 days to report, and once antibiotic treatment is established, false negative results are prone to occur (the sensitivity is approximately 78.4%)[13]. CrAg detection has a sensitivity and specificity of 99% in CSF, but latex agglutination (LA) is more expensive, more labor-intensive, and requires cold-chain shipping/storage[14]. Patients with rheumatoid factor positive, tuberculous meningitis or systemic lupus erythematosus may have a false positive reaction. The LA antigen titer of patients infected with *Trichosporon* can reach 1:1000[15]. Moreover, hemolytic samples and “hook effect” caused by high concentrations of CrAg can lead to false negative results in lateral flowassay (LFA)[16]. Histopathological examinations are complicated in preparation and staining processes, and microscopic examinations require experienced staff. Therefore, it is an urgent need to develop a rapid and sensitive diagnostic technique to complement the deficiencies of existing methods.

With the characteristics of rapid and high sensitivity, various polymerase chain reaction (PCR) technologies, such as nested PCR[17], real-time PCR[18,19], and singleplex PCR[20], have been applied to CM diagnosis. However, the extraction process of cryptococcal nucleic acid is labor intensive and cumbersome. Direct operation of samples containing *Cryptococcus* also increases the risk of exposures. Therefore, the development of integrated molecular amplification detection of cryptococcal samples is of great significance. Among the existing detection technology, nested PCR shows high specificity, but it must be carried out in two steps. Real-time PCR requires the establishment of a standard curve, internal reference and precise temperature control. All of these limit the possibility of developing integrated detection. Loop-mediated isothermal amplification (LAMP) is a good way to avoid these limitations. It is characterized by simple operation, isothermal amplification, short time required, simple equipment, and has shown promising potential in integrated detection applications.

Microfluidic chips integrated with LAMP assay for detecting bacterial meningitis, such as meningitis caused by *Neisseria meningitides*[21,22], *Streptococcus pneumoniae*[21] and *Haemophilus influenzae* type b[21], have been developed and shown high sensitivity and specificity. Recent researchs on rapid integrated detection of pathogens are mainly dedicated to any link in sample pretreatment (pathogen enrichment) or amplification method (isothermal amplification) or signal detection (colloidal gold enhanced signal)[23-26], but few solutions are available for CM detection, mainly due to the tough cell wall of *Cryptococcus* and the presence of large polysaccharide capsule (accounting for approximately 70% of the whole cellular volume) in its outer layer, which makes it difficult to achieve successful nucleic acid extraction in the chip. Various methods proposed to extract and purify DNA from *Cryptococcus* included bead rupture method, enzymatic cell wall lysis method and chemical reagents lysing method, yet the first method is difficult to implement in a functional chip due to its mechanical properties. To solve this issue, we combined the use of lyticase digestion, and thermal alkaline lysis in the microfluidic chip to achieve effective nucleic acid extraction. In addition, we used a filter membrane to capture the target and a portable UVA flashlight to read the amplified signal, thus simplifying the entire operation process.

Here we developed a LAMP-based microfluidic chip to combine rapid filter membrane-based sample concentration, nucleic acid extraction, target DNA amplification, and portable UVA flashlight signal reading of pathogens from CSF samples, which demonstrated a rapid, high-efficiency approach, and enabled “sample-to-answer” detection from real CSF samples.

## Materials and Methods

### CSF samples

Between March 2019 and November 2019, 83 clinical CSF samples that were LFA positive (CrAg titers ranging 1:1–1:20,480) were collected from CM patients of Shanghai Huashan Hospital in China. A total of 40 clinical positive-cultured CSF samples collected from patients with other CNS infections (diagnosed by culture or pathology) (Table S1). Sterile human CSF was collected from a non-infected patient who required CSF drainage due to high CSF pressure, with his consent in Shanghai Huashan Hospital. With ethical approval (accession number: KY2019 HIRB-002), all patients involved signed informed consent forms, understanding and agreeing to use their clinical samples in this study.

### Isolates

Eleven standard strains containing different serotypes, genotypes and AFLP types were donated by Professor Min Chen (Table S2). Fifty strains used for specificity verification were collected from Shanghai Huashan Hospital and confirmed by MALDI–TOF mass spectrometry (Bruker Daltonics, Bremen, Germany) and nucleic acid sequence. (Table S3).

### Standard Nucleic acid extraction

Genomic DNA from cultured microorganisms was extracted using the Yeast DNA Kit (Omega Bio-Tek, Norcross, GA). Genomic DNA from the clinical samples was extracted using the QIAamp DNA Mini Kit (QIAGEN, Hilden, Germany) with some modifications: an initial incubation with lyticase (10U) at 30 °C for 45 min, followed by AL buffer incubation at 90 °C for 10 min, was included. The concentration of DNA was evaluated using Qubit 3.0 fluorometer (Invitrogen, ThermoFisher Scientific, Malaysia).

### LAMP assay performance verification

LAMP primers were designed using Primer Explorer V4 software (http://primerexplorer.jp/e/) based on the *Cryptococcus* capsular-associated protein 10 gene (CAP10) (Table 1). The assay was run on an ABI Prism 7500 real-time PCR system (Applied Biosystems, USA). The reaction was carried out in a 25 μl reaction mixture containing 13 μl 1×ThermoPol Buffer (20 mM Tris-HCl, 50 mM KCl, 10 mM (NH4)_2_SO_4_, 2 mM MgSO_4_, 0.1% Tween 20, 6 mM MgSO_4_, 1.4 mM dNTP mixture, 5×SYBR Green), 1 μl target-specific primer mixture (1.6 μM FIP/BIP, and 0.2 μM F3/B3), 1 μl 8 U of Bst DNA polymerase and 10 μl of DNA. The mixture was incubated at 64.8 °C for 5 s, 65 °C for 55 s (collecting the fluorescent) for 45 minutes. The sterile CSF and ultra-purified water were used as negative controls. As a positive control, 1ng/μl purified *C. neoformans* DNA was used.

**Table 1.**
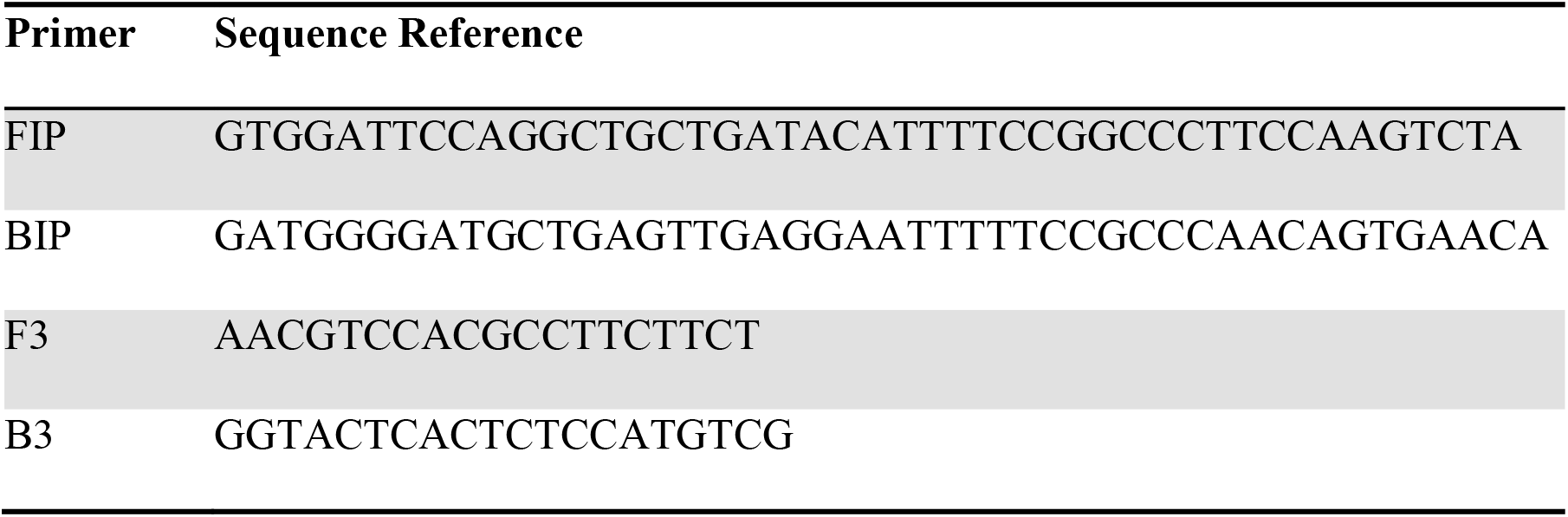
Sequence of the LAMP primers

Evaluation of the accuracy of the LAMP assay: Eleven standard strains containing different serotypes, genotypes and AFLP types were adjusted to 0.5 McFarland standard suspensions and DNA was extracted for LAMP accuracy verification.

Evaluation of the specificity of the LAMP assay: DNA extracted from forty clinical positive-cultured CSF samples collected from patients with other CNS infections (including *Enterobacteriaceae, Acinetobacter, Staphylococcus, Enterococcus, Listeria sp, Candida, Nocardia, Mycobacterium, Aspergillus, Cysticercus* and complex infection) (Table S1), one sterile human CSF sample (negative control), one ultra-filtered water (negative control), and fifty pure cultures of microorganisms (including different species and closely related species) (Table S3), was tested for LAMP specificity.

Evaluation of the detection limits of the LAMP assay: Continuously dilute (1:10) the *C. neoformans* yeast suspensions and its DNA into sterile CSF ranging from 10^8^ CFU/ml to 1 CFU/ml and 1ng/μl to 1fg/μl, then extract the DNA and use it for sensitivity verification.

Evaluation of the precision of the LAMP assay: The precision assay was divided into intra-batch and inter-batch trials. The intra-batch trial selects 20 1 pg/μl weak positive samples and 20 ultra-filtered water samples for testing together. The inter-batch trial selects one 1pg/μl weakly positive sample and one ultra-filtered water sample and tests once a day for 20 days.

Analysis of clinical samples: DNA extracted from 83 LFA-positive CSF samples was used as a template for LAMP assay.

### Chip fabrication

All the microchannels and the reaction holes were fabricated using soft lithography. The PDMS (Momentive, NY, USA) prepolymer and curing agent were mixed at a ratio of 5:1 (w:w) and 7:1 (w:w) to prepare different layers. The mixture was casted onto a silicon wafer having different patterns of SU-8 2100 (MicroChem, MA, USA) and AZ-50 (AZ Electronic Materials, Merck, USA) and cured at 80 °C, afterwards the cured PDMS layer was peeled off from the wafer. Holes in reaction layer, mixed layers and filtration layers were punched with 2mm and 5mm perforators respectively. The 1 μm pore diameter sized polycarbonate membranes (General Electric, WI, USA) were sealed between the mixed layers and cured at 80 °C for 6 h. A ratio of 10:1 PDMS prepolymer/curing agent was injected into the filling channel through the filling inlet and was cured at 120 °C for 5 min to solidify the filtration layers sandwich. Then, the reaction layer, collection layers, mixed layers and filtration layers were aligned and heated sealing together at 80 °C overnight.

### On-chip nucleic acid extraction

In order to obtain free nucleic acid in microfluidic chip, we combined lyticase, Qiagen DNA Kit AL buffer (QIAGEN, Hilden, Germany), Novagen Bugbuster Master Mix (Novagen, Merck, USA) and three additives (NaOH, SDS, urea) to select the best extraction strategy. Besides, temperature was another influencing factor.

### On-chip LAMP amplification and signal detection

After nucleic acid extraction, the LAMP reaction solution and nucleic acid were injected into the reaction wells. Polyester (PET) film (General Electric, WI, USA) sealed the top layer, and the integrated chip was incubated in an oven at 65 °C for 45 minutes. After on-chip LAMP amplifications, a portable UV flashlight was applied to shine LAMP products to obtain the visual detection results. The generated fluorescence could be observed by the naked eye.

## Results

### LAMP assay performance verification

The accuracy of the LAMP assay: Analysis of 11 different serotypes, genotypes and AFLP types of *C. neoformans/gattii* standard strains showed that accuracy was 100% (Table S2).

The specificity of the LAMP assay: The results showed that the specificity of the assay is 100%, as both the 40 clinical samples from patients with other CNS infections and 2 negative controls (0 positive/42 samples) (Table S1) gave negative results, and no cross-reaction was observed in 50 other pure cultured microorganisms (0/50) (Table S3). The detection limits of the LAMP assay: The optimized LAMP assay conditions in our system allowed detection of 100 fg/μl of *Cryptococcus* DNA (Figure 1A) and 100 CFU/ml of *Cryptococcus* yeast suspension (Figure 1B).

**Figure 1.**
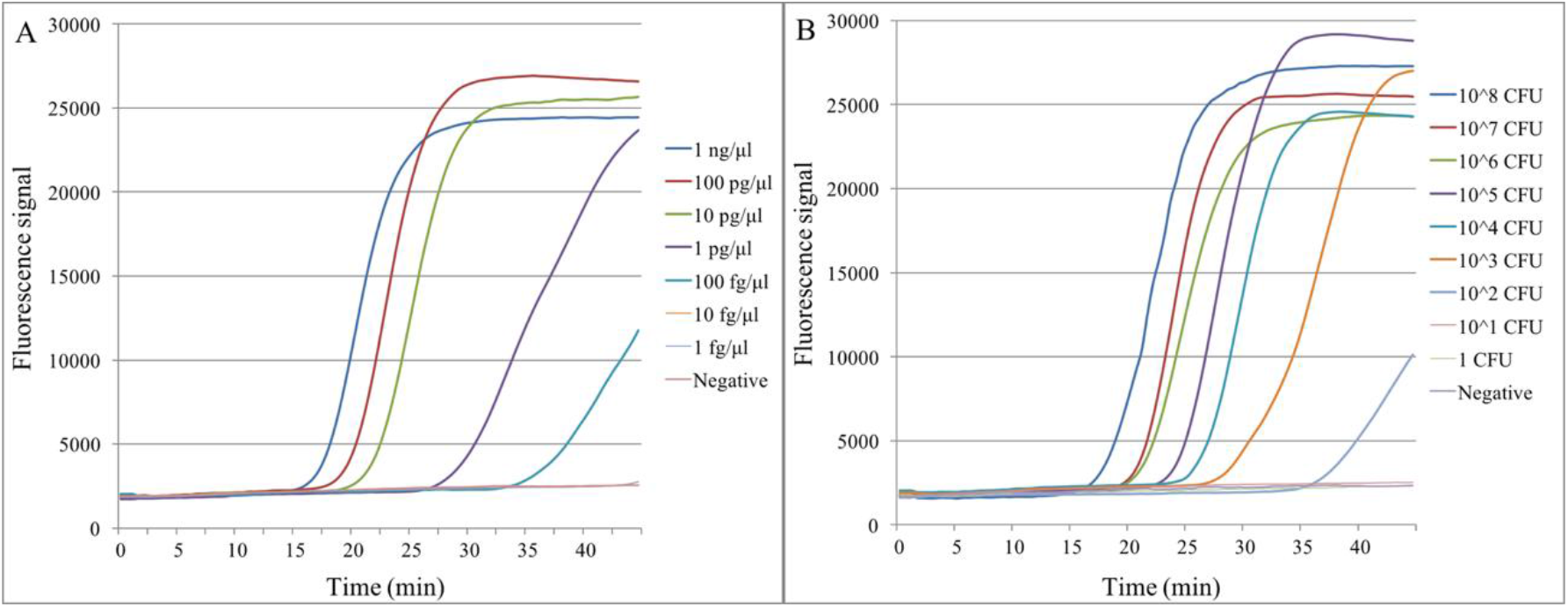
The detection limits of the LAMP assay. A: Detection limit of *Cryptococcus* DNA. The blue, brown, green, purple, light blue, yellow, light purple and pink lines represented 1ng/μl, 100pg/μl, 10pg/μl, 1pg/μl, 100fg/μl, 10fg/μl, 1fg/μl and negative templates, respectively. B: Detection limit of *Cryptococcus* yeast suspension. The blue, brown, purple, light blue, yellow, light purple, pink, light green and grey lines represented 10^8^, 10^7^, 10^6^, 10^5^, 10^4^, 10^3^, 10^2^,10, 1 CFU and negative templates, respectively.

The precision of the LAMP assay: In intra-batch and inter-batch precision tests, 20 weakly positive samples (1pg/μl) were all positive, 20 negative samples were all negative, and the agreement rates were 100%. The average positive time of the respective intra-batch and inter-batch assay was 26.82/27.33 min, SD value was 1.70/3.02, and CV value was 6.33%/11.06%.

Analysis of clinical samples: The total positive rate of 83 CSF samples detected by LAMP PCR was 91.6% (76/83). The positive rate of LAMP assay with LFA titer in the range of 1:80-1:2,040 was 100%, while those of lower titers (1:1-1:40) was 33.3-90.0% (Table 2).

**Table 2.** LAMP results for clinical samples from patients with proven CM

| Titres of CrAg | N | + | - | Positive rate (%) |
| --- | --- | --- | --- | --- |
| 1:20480 | 3 | 3 | 0 | 100% |
| 1:10240 | 8 | 8 | 0 | 100% |
| 1:5120 | 3 | 3 | 0 | 100% |
| 1:2560 | 4 | 4 | 0 | 100% |
| 1:1280 | 8 | 8 | 0 | 100% |
| 1:640 | 8 | 8 | 0 | 100% |
| 1:320 | 8 | 8 | 0 | 100% |
| 1:160 | 10 | 10 | 0 | 100% |
| 1:80 | 7 | 7 | 0 | 100% |
| 1:40 | 10 | 9 | 1 | 90.0% |
| 1:20 | 7 | 5 | 2 | 71.4% |
| 1:10 | 3 | 1 | 2 | 33.3% |
| 1:5 | 2 | 1 | 1 | 50.0% |
| 1:1 | 2 | 1 | 1 | 50.0% |
| Total | 83 | 76 | 7 | 91.6% |

### Microfluidic chip pathogen enrichment

This core area was composed of 4 duplicate filtration membrane structures (Figure2). PDMS/curing reagent mixtures were injected through the filling inlet and the protruding small dots around the corner were used as a ventilation structure to enable the mixture injection more easily. The CSF sample was injected into the inlet and evenly distributed to four enrichment zones. After washing with deionized water, pathogens were effectively enriched and purified (Figure 3).

**Figure 2.**
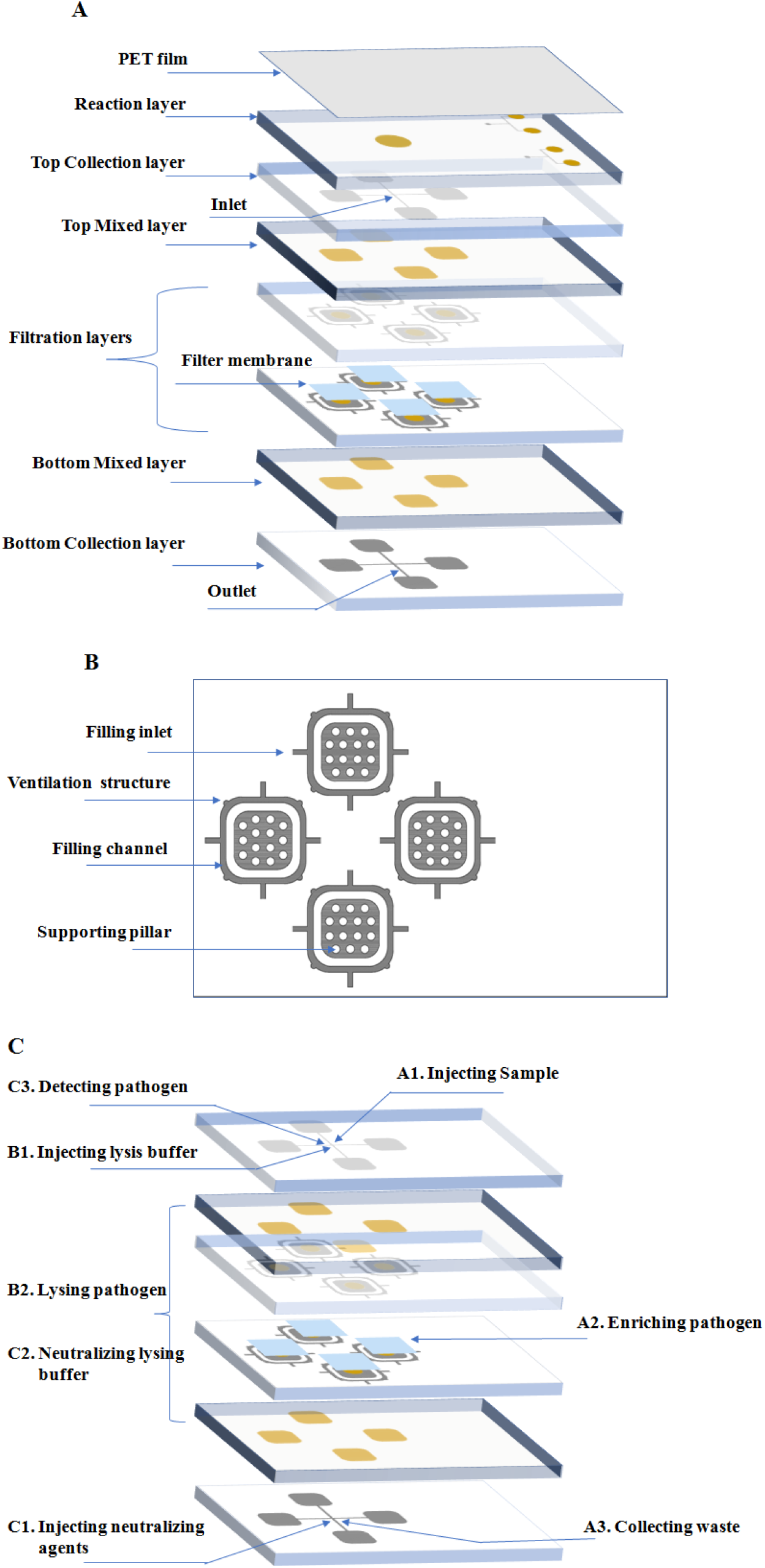
Illustration of this microfluidic chip. A) The integrated chip has nine layers, including functional layers and other different layers. The yellow part on these layers indicates that the area has been punched through, and the gray part indicates that the area is a fluid channel or a sinking part. B) This core area consisted of 4 duplicate parts. PDMS/curing reagent mixtures were injected through the filling inlet and the protruding small dots around the corner were used as a ventilation structure. Supporting pillars were used to support the filter membrane to improve filtration efficiency. C) The nucleic acid extraction process of the microfluidic chip included three steps. A. Pathogen enrichment using filter membrane; B. Pathogen lysing with optimized lysis buffer; C. Pathogen nucleic acid protection with neutralized agents.

**Figure 3.**
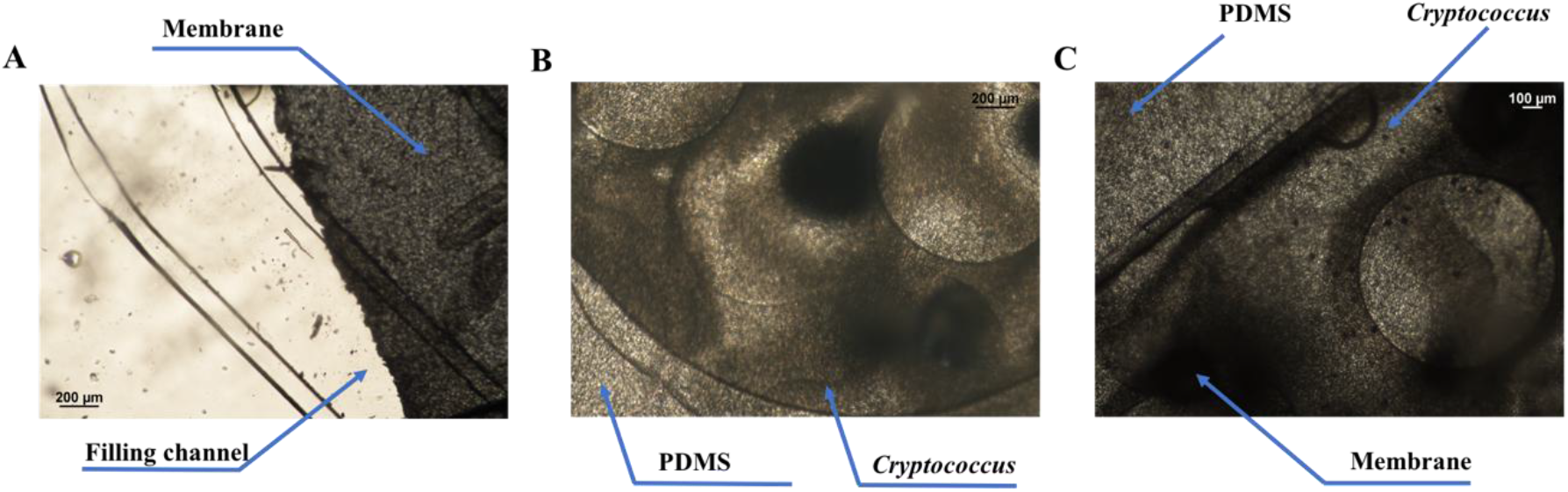
Filtration membrane enrichment images. A) Image of filtration membrane structure before *Cryptococcus* injection; B) Image of filtration membrane structure after *Cryptococcus* injection; C) *Cryptococcus* enriched by filtration membrane was stained by black ink.

### Optimization of the on-chip nucleic acid extraction strategy

Three different additives (NaOH, SDS and urea) were tested using standard Qiagen AL buffer, and it was found that urea and SDS inhibit the amplification if removed unclearly. Compared with other two additives, 0.5M NaOH improved the extraction efficiency (Figure 4A). We also compared the mild Bugbuster buffer (which was widely used to lyse bacteria and had little influence on amplification) with Qiagen AL buffer using NaOH additive and found that the lysis capacity of Bugbuster buffer was not enough for *Cryptococcus* (data not shown). Moreover, considering the tolerance of the chip temperature, we selected 25 °C, 45 °C, 65 °C and 85 °C as test points under the aforementioned conditions, and incubated the lysing mixture for 5 minutes. The amplification results showed that the lysis efficiency could be improved under incubation at 65 °C or 85 °C (Figure 4B). To simplify the process, we chose 65 °C as the lysis temperature. Then, we explored the experimental effects of different concentrations of lyticase to optimize the best working concentration. The results showed that 40U/ml lyticase digested *Cryptococcus* at 30 °C for 30 minutes in the 1M sorbitol buffer environment, and the amplification effect was the best (Figure 4C).

**Figure 4.**
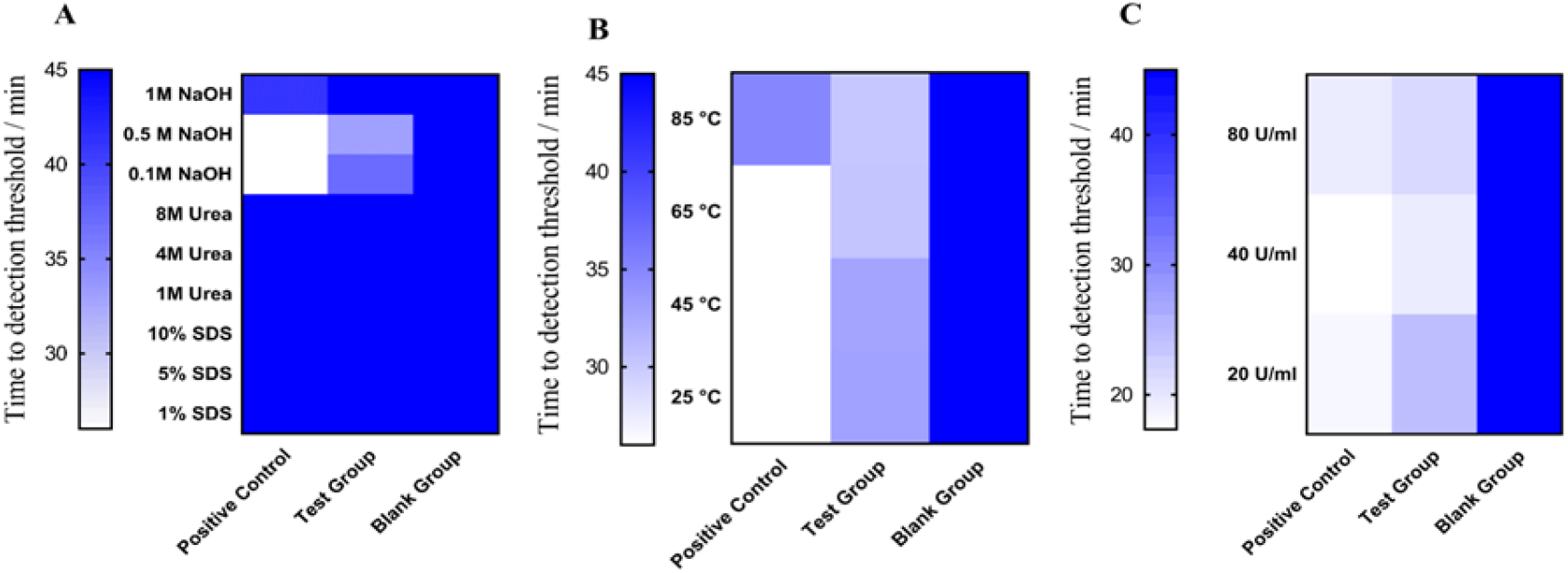
Nucleic acid fast extraction strategy. The positive control, test group and blank group, (referring to the nucleic acid extracted by the standard method, the sample and the DNase-free water respectively) were put into the lysis buffer, and then detected by the LAMP reaction after nucleic acid extraction. A) Three additives were tested for lysing capability and influence on LAMP reaction; B) Four temperature points were tested for lysing capability and influence on LAMP reaction. C) 1M Sorbitol buffer containing different concentration of lyticase was tested for its lysing capability and influence on LAMP reaction.

### Performance of the chip

The integrated chip was composed of functional layers (the reaction layer, collection layers, mixed layers and filtration layers) and other different layers. The illustration and the entire operation process were shown in Figure 2, Figure 5 and supplementary material video1. The sample was loaded into the filter well. After *Cryptococcus* enrichment, 300μl sterile deionized water was injected to wash the membrane to remove the matrix. The liquid was evacuated with air, and approximately 60μl of 1M sorbitol buffer containing 40U/ml lyticase was injected through the inlet, and withdrawn from the outlet. After 5 minutes of repeated suction operation, the filter membrane structure rich in pathogens was evenly immersed in the buffer environment, and incubated at 30 °C for 30 minutes. Subsequently, 60 μl of AL buffer containing 0.5M NaOH was injected. Repeated the above suction operation, and after the lysis buffer was in full contact with the *Cryptococcus*, incubated the chip at 65 °C for 5 minutes to release the nucleic acid. Then, 100 μl of 100 mM PH 8.0 Tris-HCl buffer was injected through the outlet, and the lysis buffer was neutralized by completely mixing from the top layer to the bottom layer to release free nucleic acids. The LAMP reaction solution and DNA were injected into the reaction well for isothermal amplification for 45 minutes. After 45 minutes of amplification,the detection results could be successfully read by the naked eye (Figure 6).

**Figure 5.**
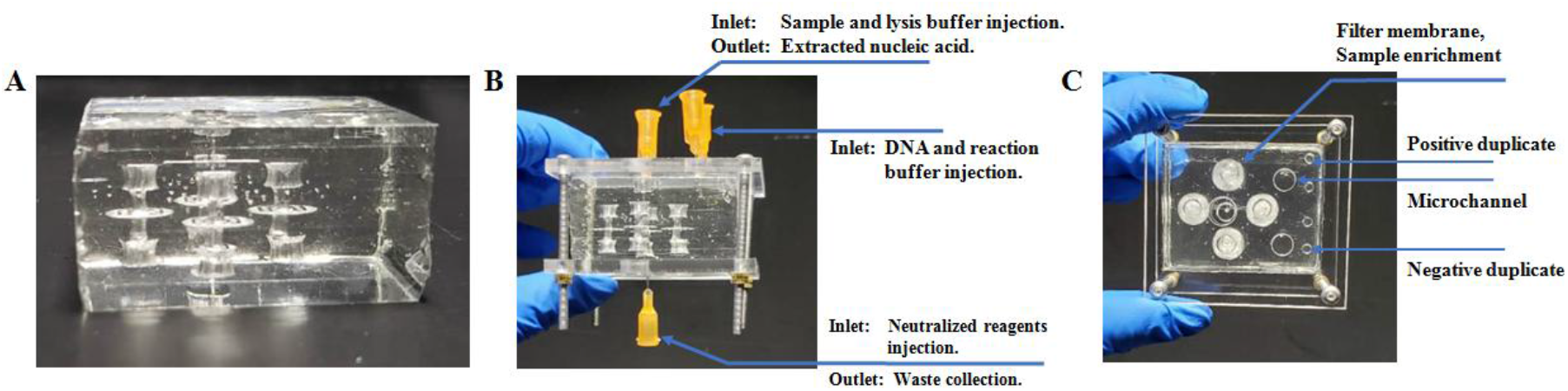
Picture of integrated chip. A) Integrated chip with PET film sealed; B) Integrated chip inlet and outlet; C) Integrated chip top view, including filter membrane, detection well, inlet, outlet and microchannels.

**Figure 6.**
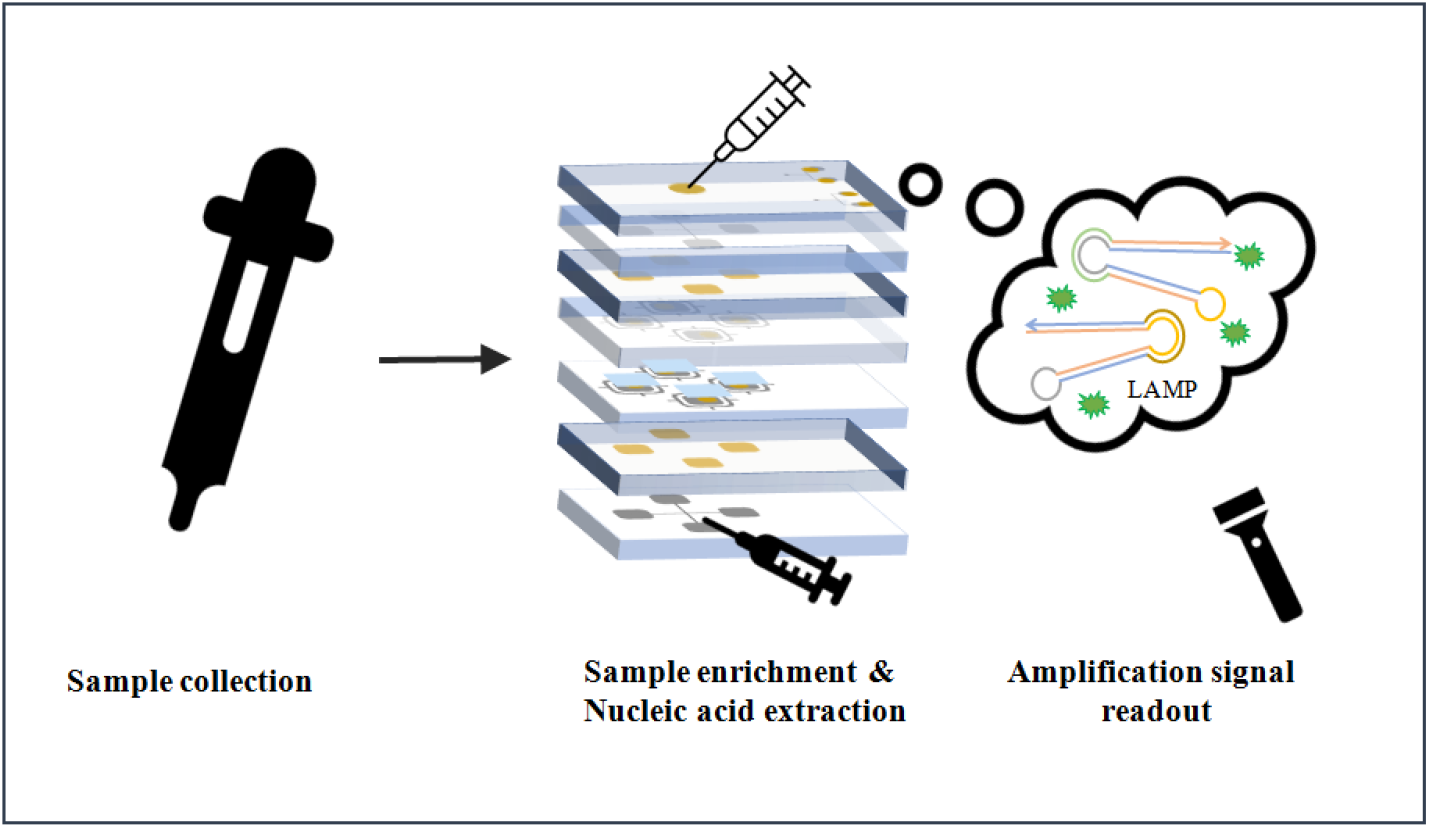
The whole process of detecting *Cryptococcus* on microfluidic chip.

## Discussion

Cryptococcal meningitis is a global threat with significant attributable mortality. Worldwide, CM is typically associated with HIV infection[2], and it is increasingly recognized in patients without HIV[6-11]. To date, information about integrated detection for CM diagnosis is still limited. In this study, we developed a LAMP-based microfluidic chip for the diagnosis of CM.

In order to ensure the accuracy and reliability of the LAMP system, we conducted LAMP assay performance verification. We collected 11 different serotypes, genotypes and AFLP types of *C. neoformans/gattii* standard strains for accuracy verification, which was 100% and showed its high accuracy. The agreement rates of intra-batch and inter-batch precision tests were 100%. The average positive time of the intra-batch and intra-batch assay was respectively 26.82/27.33 min, SD value was 1.70/3.02, and CV value was 6.33%/11.06%, which showed relatively stable reproducibility.

Generally, internal transcribed spacers have been used for molecular identification of fungal, but unfortunately, these sequences have little discrimination between *C. neoformans*/*gattii* and their closely related species, impairing their use in species differentiation by PCR methods[27]. To avoid non-specific amplification, we chose CAP10 gene to design LAMP primers, mainly considering that it encoded a specific capsular protein of *C. neoformans*/*gattii*, with little homology to their related species. The results showed the specificity was 100%, demonstrating the high specificity of the LAMP systems that we established.

The sensitivity of the LAMP assay for *Cryptococcus* cells is 100 CFU/ml. Significantly, it is of great importance for persons presenting early in the CM process with lower burden of infection, because India ink’s sensitivity was only 42% when the *Cryptococcus* CFU value is <1,000 per ml of CSF[13]. Although the detection limit of LFA also reached 100 CFU/ml, LAMP assay could confirm cases which LFA had questionable results. Moreover, the sensitivity of the LAMP assay for *Cryptococcus* genomic DNA was 100 fg/μl, which was little lower than those of Min Chen *et al*. (20 fg genomic DNA tested by LAMP assay with turbidity method)[28] and Sara Gago *et al*. (2 fg genomic DNA tested by Real-time PCR)[18,19]. The potential reason may be that we added the cryptococcal DNA to the sterile CSF for further DNA extraction, which consumed a portion of the DNA, while they used DNA to detect directly. Accordingly, we determined the extraction efficiency of approximately 53.5%. Therefore, the detection limit of cryptococcal DNA in our study was similar to that of the Min Chen *et al*. study[28].

In clinical specimen verification, the total positive rate of 83 CSF samples detected by LAMP PCR was 91.6% (76/83), which was higher than previously reported LAMP assay with turbidity method (87.1%, 74/85)[28] or real-time PCR (90.7%, 39/43) results[19]. Although the positive rate of LFA titer in the range of 1:80-1:2,040 was 100%, those of lower titer (1:1-1:40) was 33.3-90.0%. The potential reason might be that dead *Cryptococcus* cells continue to release capsular polysaccharide antigen, and the body clears the antigen relatively slowly. Even after several months of effective treatment, the results of LFA could still be positive. Therefore, the results of a patient with low LFA titer could not truly reflect the burden of *Cryptococcus* in vivo.

In all, we established a rapid, accurate, sensitive, specific, and reproducible LAMP assay to detect *C. neoformans/gattii* in CSF samples. This assay might be an alternative method for rapid diagnosis of cryptococcal meningitis, especially for those with low cryptococcal load. Moreover, our research confirmed the feasibility of this LAMP system, which made a good foundation for the further development of microfluidic chip.

Developing a microfluidic chip for real-life application of CM diagnosis remains a challenge. Major limiting factors are low-concentration pathogen, complex sample matrices and difficulties in extraction of *Cryptococcus* DNA on the chip. For removing interfering substances and enrichment of pathogens, commonly used pretreatment tools are centrifugation, filtration membranes or immune-based techniques [29]. However, coupling the centrifugal device to the microfluidic chip is relatively complicated, and the activity of immune-based methods is unstable. In contrast, the filter membrane has a simple structure and can quickly and efficiently enrich pathogens at a level of ∼10^−1^μm (*Cryptococcus* was nearly 5μm in diameter)[30]. Benefit from these features, the filter membrane structure was conveniently integrated into our chip, thus simplifying the entire capture process. The core area of the microfluidic chip was composed of 4 duplicate filtration membrane structures, aiming to expand the filtration area, improve fluid throughput and prevent enrichment oversaturation caused by a single membrane structure. Supporting pillars were used to support the filter membrane to improve filtration efficiency. Pathogens in CSF samples were enriched through filtration membrane, washed and purified, thus improving the system sensitivity and avoiding the inconsistency of results due to the heterogeneity of sample matrix. In addition, the successful extraction of on-chip cryptococcal nucleic acid is another key issue of this microfluidic chip. Traditional methods often adopted beads mechanical capture method or enzymatic cell wall lysis method. However, limited by the characteristics of the chip, the mechanical method could not be applied. Accordingly, various experiments were carried out to optimize on-chip *Cryptococcus* nucleic acid extraction. It was confirmed that the use of standard Qiagen AL buffer and 0.5M NaOH and incubated the lysing mixture at 65 °C for 5 minutes can improve the efficiency of nucleic acid extraction in the chip. However, the above conditions were still insufficient. Because *Cryptococcus* is different from other fungi, in addition to the common cell wall, its outer layer is surrounded by a thick low electron density mucus capsule. Under the double protection of the capsule and cell wall, *Cryptococcus*is much tougher and genomic DNA extraction is more difficult. The removal of external structures to form protoplasts is essential for nucleic acid extraction. Lyticase is a fungal cell wall lysis enzyme. We explored the experimental effects of different concentrations of lyticase to optimize the best working concentration. Moreover, protoplasts were very sensitive to external conditions and prone to membrane rupture with changes in osmotic pressure. We used sorbitol buffer as the osmotic pressure stabilizer to ensure a high protoplast formation rate. The results showed that 40U/ml lyticase digested *Cryptococcus* at 30 °C for 30 minutes in the 1M sorbitol buffer environment, and the amplification effect was the best. The nucleic acid extract was amplified isothermally for 45 minutes and the detection results could be successfully read by the naked eye.

This microfluidic chip, integrating sample *Cryptococcus* enrichment, optimized nucleic acid extraction and LAMP detection unit, streamlined reaction processes and reduced the exposure risk of directly handling cryptococcal samples. It did not require any additional instruments and provided a rapid, reliable, as well as high-efficiency approach. We believed that this integrated chip truly realized the “sample-to-answer” application and could be easily used for clinical cryptococcal prediagnosis.

## Acknowledgements

We thank Professor Min Chen (Department of Dermatology, Shanghai Key Laboratory of Molecular Medical Mycology, Shanghai Institute of Medical Mycology, Changzheng Hospital, Second Military Medical University, Shanghai, China) for giving us the standard strain.

---

**Table S1.**
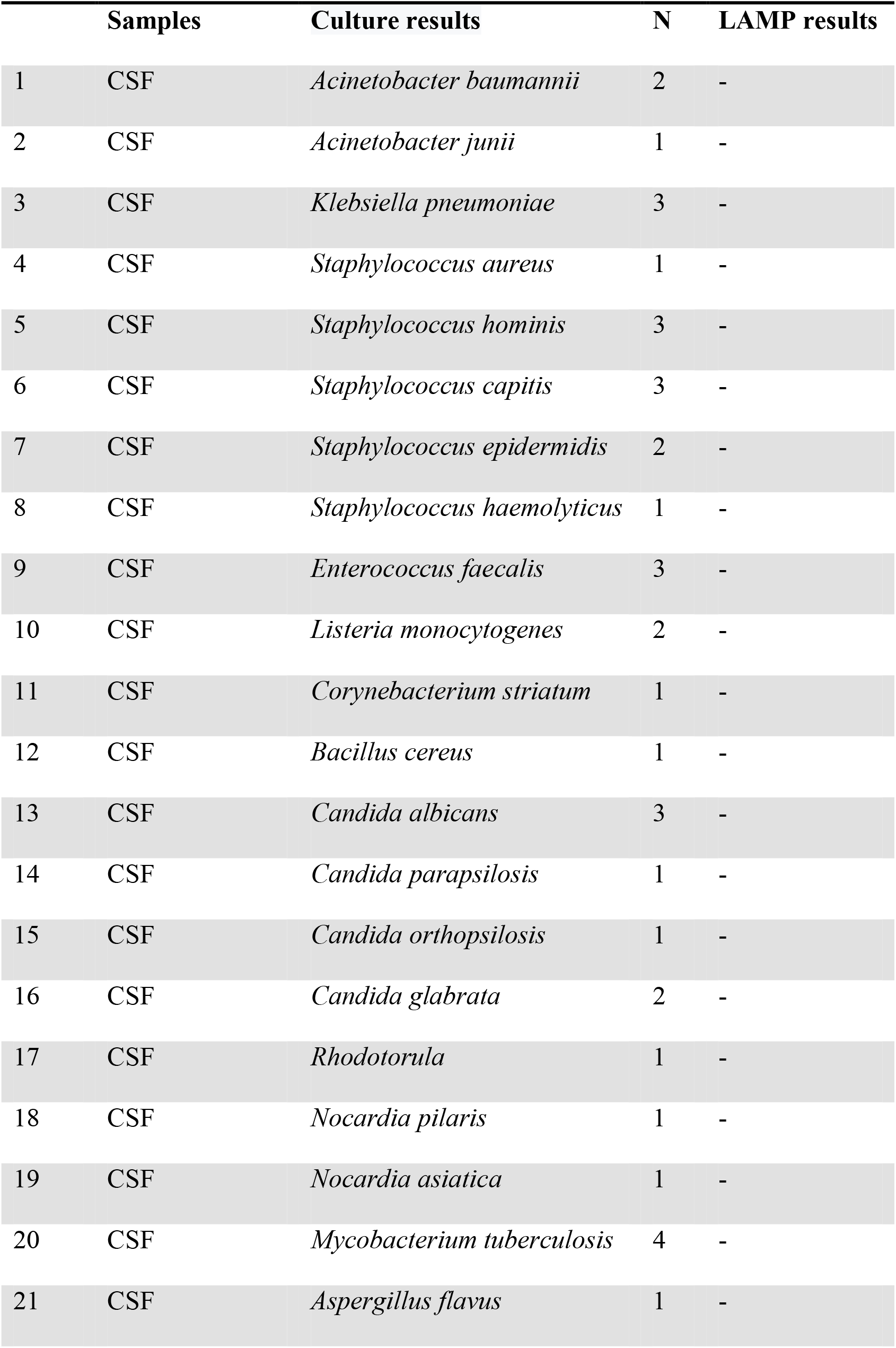
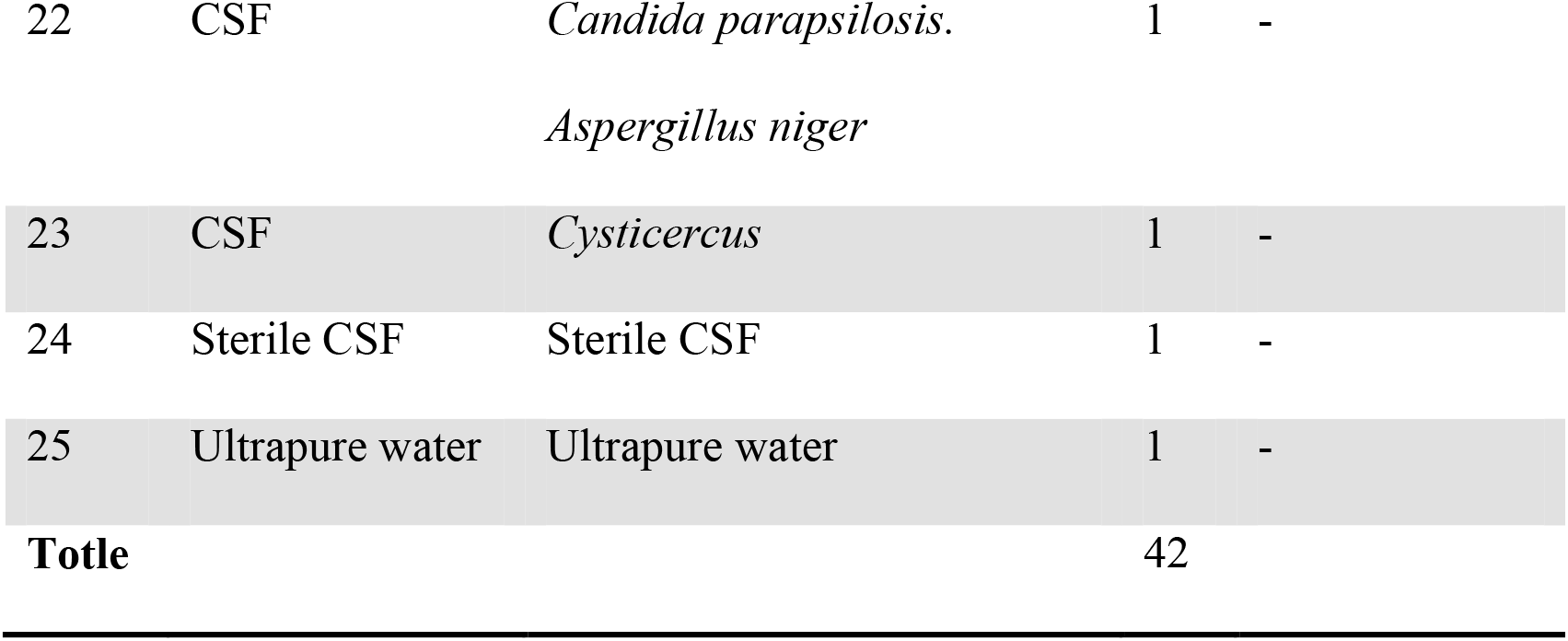
LAMP results of CSF positive samples with other CNS infections

**Table S2.** Standard *C. neoformans/gattii* strains

|  | **Standard strains** | **Serotype** | **Genotype** | **AFLP Type** | **LAMP** |
| --- | --- | --- | --- | --- | --- |
| 1 | *C. neoformans var grubii* | A | VNI | 1 | + |
| 2 | *C. neoformans var grubii* | A | VNII | 1A | + |
| 3 | *C. neoformans var neoformans* | AD | VNIII | 1B | + |
| 4 | *C. neoformans var neoformans* | D | VNIV | 2 | + |
| 5 | *C. neoformans var gattii* | B | VGI | 4 | + |
| 6 | *C. neoformans var gattii* | B | VGII | 6 | + |
| 7 | *C. neoformans var gattii* | B | VGIII | 5 | + |
| 8 | *C. neoformans var gattii* | C | VGIV | 7 | + |
| 9 | *C. neoformans var grubii* | A | VNI | 1 | + |
| 10 | *C. neoformans var neoformans* | D | VNIV | 2 | + |
| 11 | *C. neoformans var grubii* | A | VNI | 1 | + |

**Table S3.**
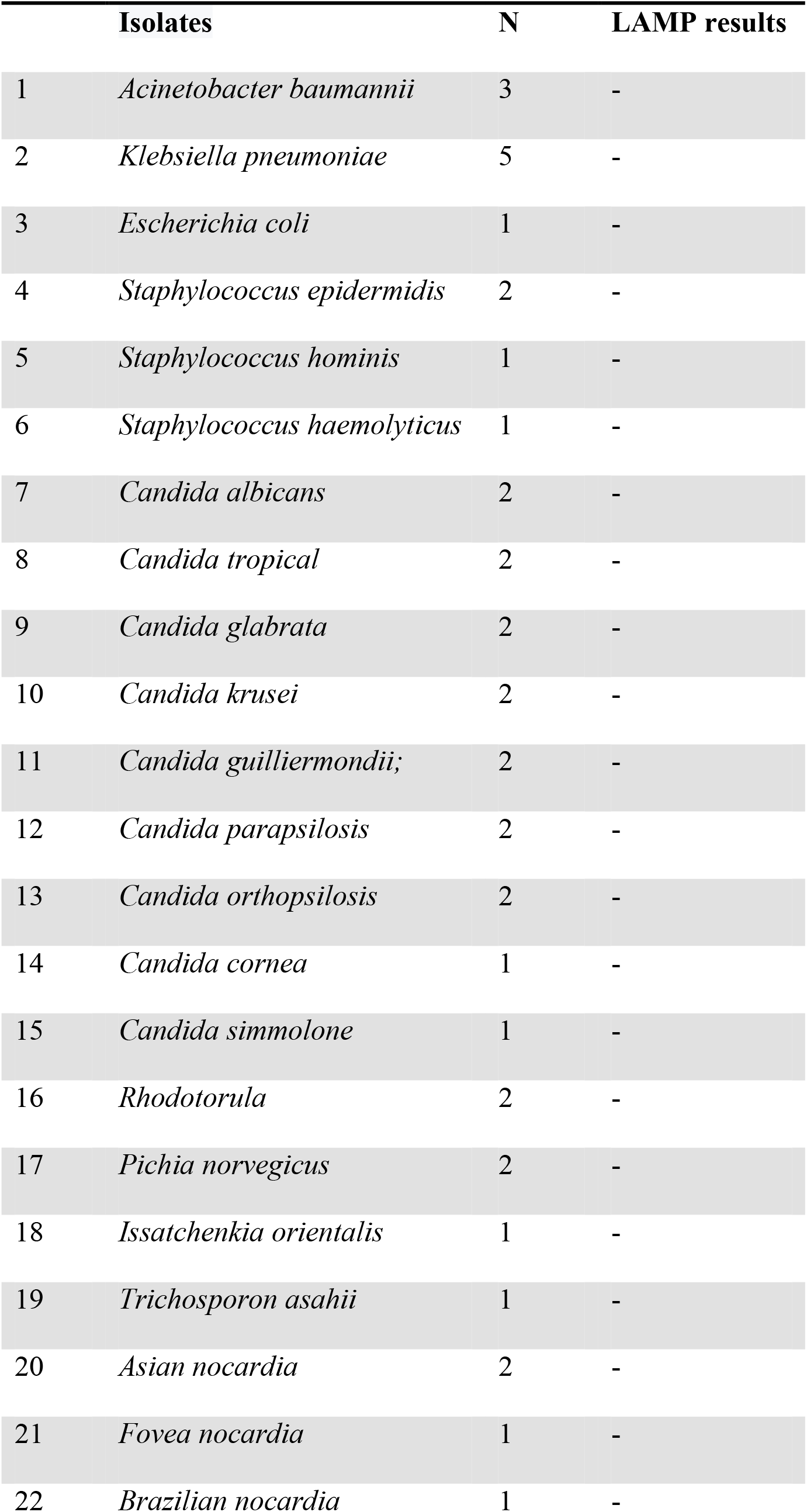
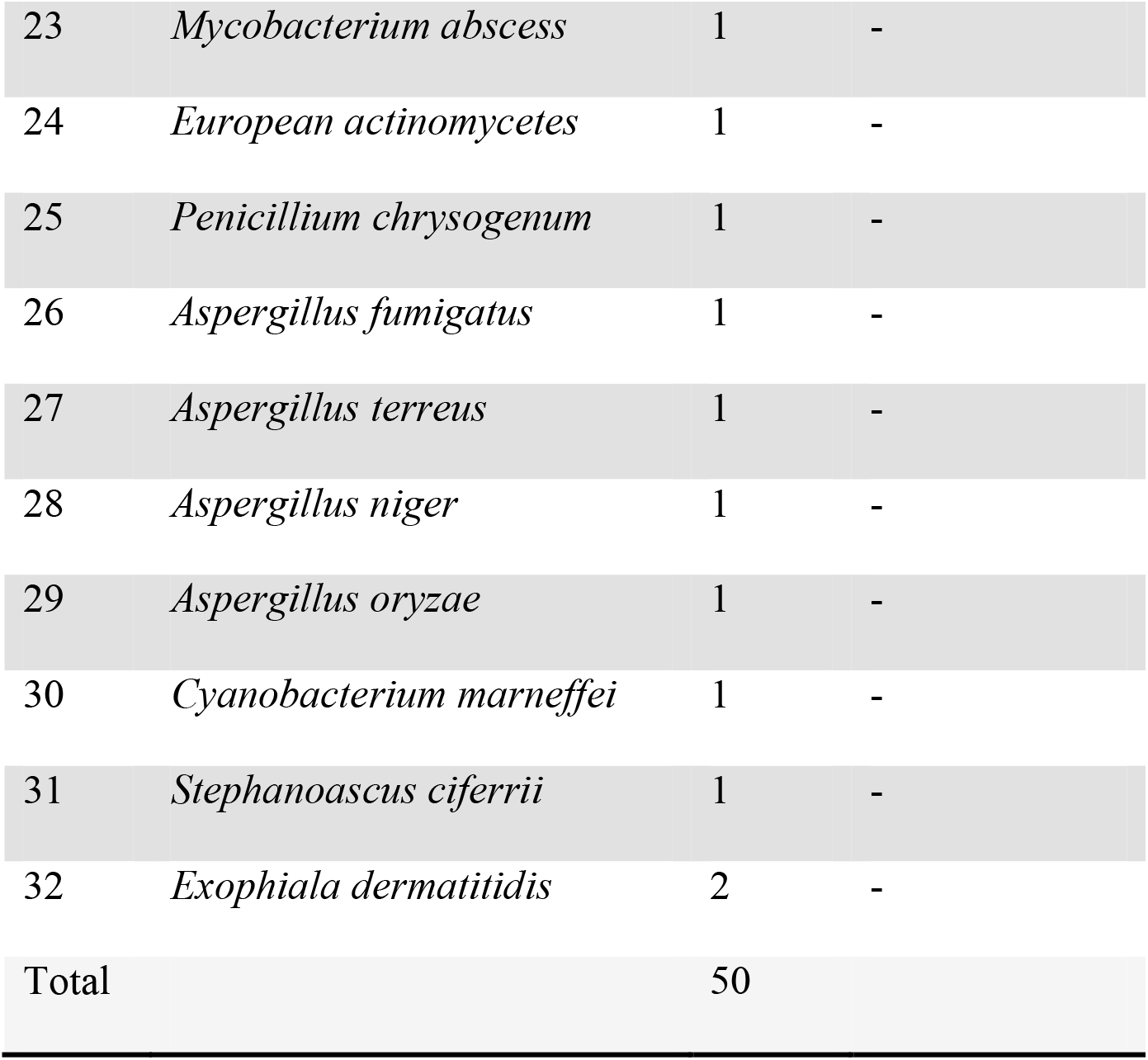
LAMP results of pure cultured isolates from different microorganisms

